# Oscillatory dynamics of Rac1 activity in *Dictyostelium discoideum* amoebae

**DOI:** 10.1101/2024.03.28.587124

**Authors:** Marko Šoštar, Maja Marinović, Vedrana Filić, Nenad Pavin, Igor Weber

## Abstract

Small GTPases of the Rho family play a central role in the regulation of cell motility by controlling the remodeling of the actin cytoskeleton. In the amoeboid cells of *Dictyostelium discoideum*, the active form of the Rho GTPase Rac1 regulates actin polymerases at the leading edge and actin filament bundling proteins at the posterior cortex of polarized cells. However, constitutive Rac1 dynamics in *D. discoideum* have not yet been systematically investigated. Therefore, we monitored the spatiotemporal dynamics of Rac1 activity in vegetative amoebae using a specific fluorescent probe. We observed that plasma membrane domains enriched in active Rac1 not only exhibited stable polarization, but also showed rotations and oscillations. To simulate the observed dynamics, we developed a mass-conserving reaction-diffusion model based on the circulation of Rac1 between the membrane and the cytoplasm in conjunction with its activation by GEFs, deactivation by GAPs and interaction with the Rac1 effector DGAP1. Our theoretical model accurately reproduced the experimentally observed dynamic patterns, including the predominant anti-correlation between active Rac1 and DGAP1. Significantly, the model predicted a new colocalization regime of these two proteins in polarized cells, which we confirmed experimentally. In summary, our results improve the understanding of Rac1 dynamics and reveal how the occurrence and transitions between different regimes depend on biochemical reaction rates, protein levels and cell size. This study not only expands our knowledge of the behavior of small GTPases in *D. discoideum* amoebae, but also provides a simple modeling framework that can be adapted to study similar dynamics in other cell types.

## Introduction

Cell migration controlled by the cortical actin cytoskeleton is the basis for important biological processes such as embryonic morphogenesis, immune surveillance, wound healing and tumor metastasis. In highly motile cells such as neutrophils, phagocytes and *Dictyostelium discoideum* amoebae, local functional domains within the actin cytoskeleton are rapidly and continuously assembled and disassembled [1,2]. These processes are coordinated spatially and temporally by complex upstream signaling pathways. Understanding these signaling networks requires studying the dynamics *in vivo* of their components, especially the small GTP hydrolases from the Rho family, such as Rac, Rho and Cdc42, which are widely recognized as important regulators of the actin cytoskeleton [3].

As molecular switches, the Rho GTPases cycle between an active GTP-bound state and an inactive GDP-bound state. Activation is facilitated by guanine nucleotide exchange factors (GEFs), and inactivation by GTPase-activating proteins (GAPs). In their active state, Rho GTPases bind and activate effector proteins, including actin-binding proteins, actin polymerases, and their direct regulators. For example, Rac interacts with formins and the Scar/WAVE complex, while Rho regulates myosin II-driven contractility [4–6]. *D. discoideum* cells, which share Rho GTPase-dependent signaling mechanisms with mammalian cells, are well suited for the study of these pathways [7,8]. This is particularly true for the three Rac1 isoforms of *D. discoideum*, which show 90% sequence identity with human Rac1 and have a 100% identical nucleotide-binding domain [7]. In *D. discoideum* cells, Rac1 interacts with proteins that play a role in shaping and regulating different structural regions of the actin cortex [7,9]. At the protruding leading edge of polarized motile cells and at the outer edge of circumferential protrusions associated with macroendocytosis, Rac1-GTP activates WASP family proteins and promotes Arp2/3-mediated branched actin polymerization [10]. The non-protruding regions of the cellular cortex are stabilized by proteins that cross-link and bundle actin filaments, such as cortexillins I and II [11]. The bundling ability of cortexillins depends on the formation of a complex with the IQGAP-related protein DGAP1 - a process catalyzed by activated Rac1 [12]. Rac1 is therefore of central importance in controlling antagonistic processes that lead to the formation of different actin structures [13,14]. In addition to the WASP and IQGAP protein families, Rac1 has been shown to interact with various other effectors, such as formins, coronins, filamins, PAK kinases and other proteins involved in actin cytoskeleton remodeling [7].

Various imaging techniques have been used to study Rho-GTPase activities and their effects on cell polarization, morphology and migration. Most probes for active GTPases are based on fluorescently labelled GTPase binding domains (GBDs) of effector proteins, either alone or incorporated into FRET constructs that detect GTPase activation by RhoGEFs [15–17]. In some experiments, multiple probes were introduced into single cells to observe spatial and temporal correlations between Rac, Rho and Cdc42 activities within protruding lamellipodia [18]. In addition to the “passive” probes that effectively reflect the activity of endogenous GTPases, there are also molecular tools developed for their selective photoactivation, mainly by manipulating associated RhoGEFs [19–22]. In summary, these methods have elucidated the intricate interplay between Rho GTPases and other proteins in actin-driven processes in a range of experimental models and conditions [23,24].

Theoretical modelling has long complemented experimental work to understand the intracellular dynamics of Rho GTPases [25]. In both budding and fission yeast, activated Cdc42 induces stable polarization by promoting actin polymerization and cell outgrowth [26]. Reaction-diffusion models have successfully simulated this polarization by incorporating a positive feedback loop between the activity of Cdc42 and the recruitment of its activating GEF, Cdc24 - a mechanism strongly supported by experimental evidence [27,28]. Similar models using mass-conserving and wave-pinning approaches simulated stable polarization of Cdc42 and Rac1 in neutrophils and macrophages [29,30]. Models with GEF-mediated negative feedback have predicted bipolar and oscillatory Cdc42 distributions, in contrast to unipolar patterns in models with only positive feedback [31].

In motile *D. discoideum* cells, actin and associated signaling proteins spontaneously form traveling waves at the ventral cell border [32]. Analyzing these patterns, as well as patterns induced by external stimuli, can facilitate the understanding of complex signaling networks and form the basis for the development of advanced theoretical models [33,34]. In this study, we show that active Rac1 exhibits a variety of dynamic behaviors in *D. discoideum* cell plasma membrane. In particular, we observed that Rac1 and its effector DGAP1 not only spontaneously form oscillating and rotating patterns, but also switch between these dynamic patterns and stationary or irregular distributions over time. To explain these observations, we constructed a reaction-diffusion model centered on the membrane binding and release of Rac1 mediated by the formation of complexes with the cognate GAP and with DGAP1. Our model faithfully reproduced the experimentally observed dynamics of Rac1 and DGAP1 as well as the correlations between the spatiotemporal patterns of the two proteins. These results suggest that the shuttling of Rac1 between the membrane and the cytoplasm, which is controlled by its interactions with regulatory proteins, adequately explains the observed oscillations.

## Results

### Rac1-GTP shows a repertoire of regular dynamic patterns

To characterize the dynamics of Rac1 activity in *D. discoideum*, we used confocal microscopy to observe vegetative cells expressing the recently developed fluorescent probe DPAKa(GBD)-DYFP, which specifically binds to Rac1-GTP [35]. For simplicity, we refer to the fluorescent signal corresponding to Rac1-GTP labeled with DPAKa(GBD)-DYFP as Rac1* in the following text. Our observations revealed that Rac1* is mainly localized at the membrane and forms patterns that are either random or ordered and usually persist for several minutes. Of the 85 cells expressing DPAKa(GBD)-DYFP that we observed, 26 showed ordered Rac1* patterns, while the patterns in the remaining 59 cells were classified as random. The ordered patterns can be categorized into three main types: Traveling waves (rotations), standing waves (oscillations) and stationary inhomogeneous distributions (polarizations). Each pattern type has either a single Rac1*-enriched membrane domain or two such domains. Based on this criterion, we further classified the observed patterns into six distinct categories: rotating monopoles (n=14, Fig 1A, S1 Movie), rotating dipoles (n=3, S1 Fig A, S2 Movie), oscillating monopoles (n=5, Fig 1B, S3 Movie), oscillating dipoles (n=2, S1 Fig B, S4 Movie), stationary monopoles (n=4, Fig 1C, S5 Movie) and stationary dipoles (n=1, S1 Fig C, S6 Movie).

**Fig 1.**
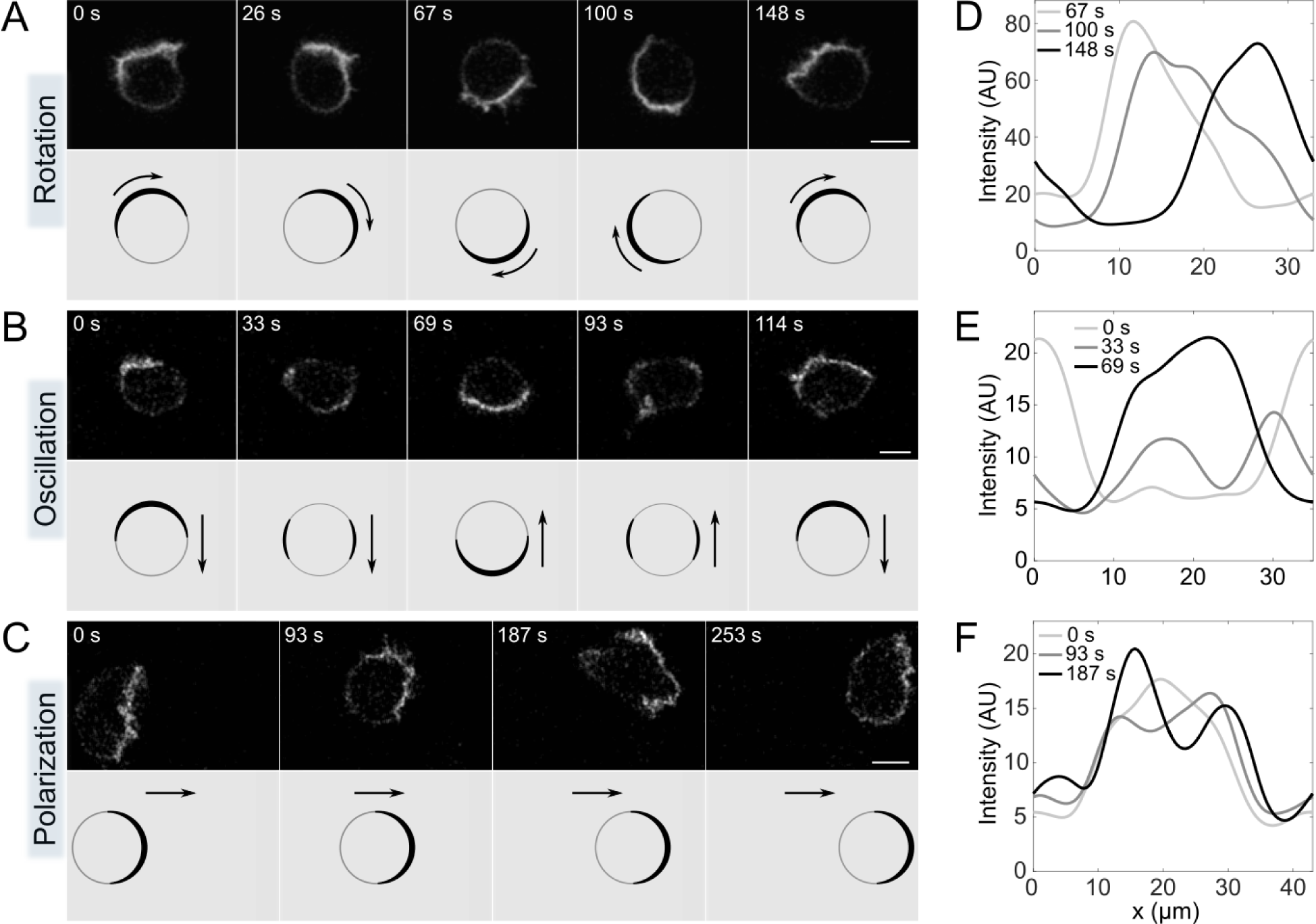
Representative monopolar patterns of Rac1*-enriched cortical domains. (**A**) *Rotating monopole.* Selected frames show the rotation of a Rac1*-enriched membrane domain (top). The corresponding drawing shows the direction of the rotating domain, as indicated by arrows (bottom). (**B**) *Oscillating monopole.* Selected frames show the periodic movement of the Rac1* domain from one side to the opposite side of the cell membrane (top). The corresponding drawing shows the direction of movement of the active domain, marked by arrows (bottom). (**C**) *Stationary monopole.* Selected frames show a Rac1* domain that remains stationary at the front of a stably polarized migrating cell for several minutes (top). The corresponding drawing shows the direction of cell migration indicated by arrows (bottom). (**D-F**) Diagrams showing Rac1* intensity as a function of position on the membrane at three different time points, represented by different shades of gray: (**D**) *Rotating monopole*; (**E**) *Oscillating monopole*; (**F**) *Stationary monopole*. Scale bars: 5 μm.

A typical Rac1* traveling wave manifests as a peak distribution that moves along the membrane at a constant speed and has an approximately constant amplitude (Fig 1D). We determined the average cycle duration of these traveling waves to be 190±40 s (mean±std). Conversely, a typical standing wave is characterized by the periodic transfer of Rac1* from one side of the cell to the other. During this transfer, a portion of Rac1* moves along the membrane in two separate sections moving in opposite directions, resulting in an intermediate state characterized by two maxima (Fig 1E, time point 33 s). The average cycle duration of the standing waves was 140±20 s. In polarized cells, the Rac1*-enriched membrane domain maintains its position for several minutes, although the intensity profiles in this domain fluctuate on shorter time scales (Fig 1F).

### Rac1-GTP and DGAP1 show similar dynamic patterns that are predominantly anti-correlated

It has been shown that the activated form of Rac1 binds specifically and directly to DGAP1, establishing DGAP1 as a Rac1 effector [12]. To compare the dynamics of Rac1 and DGAP1, we observed cells expressing both DPAKa(GBD)-DYFP and DGAP1-mRFP, which we refer to as DGAP1* in the remainder of the text. We found that both probes showed rotations (Fig 2A, S7 Movie), oscillations (S2 Fig, S8 Movie) and stable polarizations (Fig 2B, S9 Movie). The patterns of Rac1* and DGAP1* were predominantly anti-correlated, which is consistent with our previous results [13,14]. However, we recorded a few cases in which the signals from both probes were positively correlated and showed simultaneous enrichment at the leading edge of the polarized cells (Fig 2C, S10 Movie). The data show that of the 93 cells observed expressing both probes, 34 cells exhibited ordered patterns, while 59 cells showed random behavior. We observed 20 rotations (12 monopoles and 8 dipoles), 9 oscillations (6 monopoles and 3 dipoles), 6 polarized cells in which the two probes were localized at opposite poles (4 monopoles and 2 dipoles), and 2 cells in which the probes were co-localized (1 monopole and 1 dipole). The average period of the travelling waves was 200±50 s, and that of the standing waves was 160±20 s. Compared to the dynamics of DPAKa(GBD)-DYFP expressed alone, these results also show that the expression of DGAP1-mRFP does not significantly affect the dynamics of Rac1*. Taken together, our experiments reveal generic, regular patterns in the membrane distribution of active Rac1 in unstimulated vegetative *D. discoideum* cells and their close relationship to the corresponding, predominantly anti-correlated patterns of the Rac1 effector DGAP1. Regular patterns are particularly useful to uncover the processes governing Rac1 dynamics, as they depend on the underlying biochemical reactions. We therefore decided to use these ordered patterns as a basis for investigating the key processes involved in the regulation of Rac1 dynamics through computational modelling.

**Fig 2.**
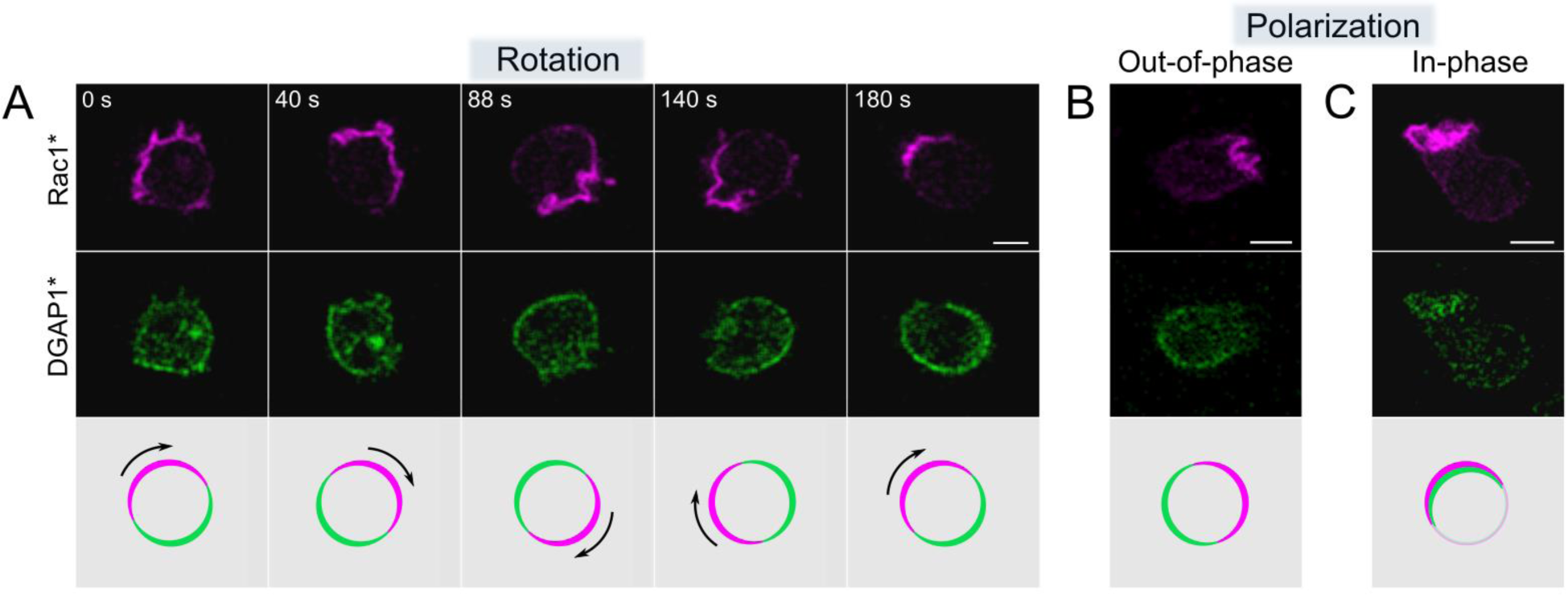
Representative patterns of cortical domains enriched in Rac1* and DGAP1*. (**A**) *Rotating monopole.* An image sequence showing a complete rotation of domains rich in Rac1* and DGAP1*. Top: Rac1* signal. Centre: DGAP1* signal. Bottom: a superposition of both fluorescent domains. In areas where Rac1* predominates, the DGAP1* signal is attenuated and vice versa. (**B**) *Stationary monopole with opposite localization.* Rac1* and DGAP1* are enriched at opposite positions. The image shown is representative of a sequence spanning 170 s. (**C**) *Stationary monopole with co-localization.* Rac1* and DGAP1* signals are co-localized. The image shown is representative of a sequence extending over 80 s. Scale bars: 5 μm.

### Computational model of Rac1 dynamics that takes into account the interaction with its effector DGAP1

To explain the mechanisms underlying the observed spatio-temporal distributions of Rac1* and DGAP1*, we propose a simplified model based on current knowledge of the interactions between Rho GTPases and their regulators and effectors. The reactions involved are graphically represented as two cycles (cycle i and ii) to visually separate the reactions of Rac1-GTP (Rac1_T_ for short) with its regulator GAP from the reactions with its effector DGAP1 (Fig 3A). A detailed discussion of the individual reaction terms included in the model can be found in Appendix 1. Both cycles in our proposed model involve the binding step of Rac1-GDP (Rac1_D_ for short) to the membrane, coupled with its conversion to Rac1_T_ through the action of a GEF [24,36,37]. In our model, this process also involves autocatalytic GEF-mediated activation of Rac1, which has been experimentally demonstrated for Rho GTPases in various cellular systems and is an established component of most theoretical models of Rho-GTPase dynamics [26,38,39]. In *D. discoideum*, RacGEF1 has also been shown to be involved in a positive feedback loop that promotes Rac1 activity [40].

**Fig 3.**
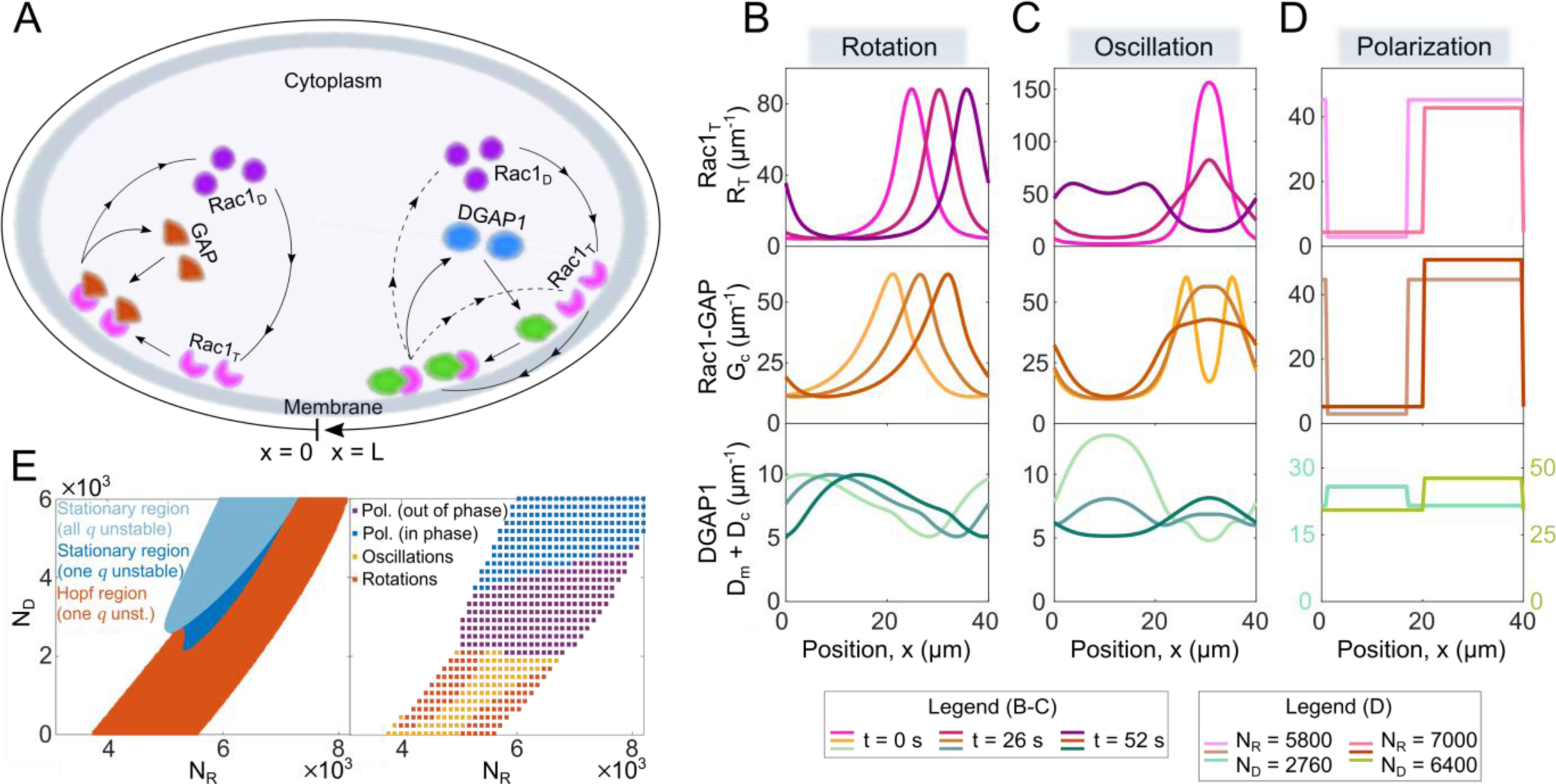
Model of the interaction network and generated patterns. (**A**) Schematic representation of the protein interaction network. Cycle (i) is shown on the left. Cytoplasmic Rac1_D_ (represented by purple disks) converts to its active form, Rac1_T_, immediately after binding to the membrane. GAP proteins (shown as brown triangles) can bind to active Rac1, leading to its deactivation and the breakdown of the Rac1_T_-GAP complex, resulting in GAP and Rac1 returning to the cytoplasm. Cycle (ii) is shown on the right. DGAP1 proteins from the cytoplasm (blue ovals) bind to the membrane where they form a complex with Rac1_T_. When the Rac1_T_-DGAP1 complex dissolves, DGAP1 moves back into the cytoplasm, while Rac1 is either released into the cytoplasm or remains attached to the membrane. (**B-D**) Traveling waves, standing waves and polarized patterns of membrane-bound proteins obtained from simulations. For traveling and standing waves, the plots show the molecular density versus position in space for three different time points. For the polarized patterns, the densities are shown for two different total copy numbers of Rac1 and DGAP1 proteins. (**E**) *Left* Diagram showing different types of instabilities as a function of the total number of Rac1 and DGAP1 proteins, as determined by linear stability analysis. In the uncolored areas of the diagram, small perturbations decay and the system reaches a homogeneous steady state. *Right* Diagram of the different dynamic states as a function of the total number of Rac1 and DGAP1 molecules, as determined by examining the final pattern produced by the system in computer simulations. The uncolored sections of the diagram correspond to the homogeneous states. All parameter values are listed in S2 Appendix Table 1.

Cycle (i) further describes the binding of cytoplasmic RhoGAP to membrane-bound Rac1_T_, which promotes its conversion to Rac1_D_, coupled with the release of both proteins into the cytoplasm. Here we assume that only Rac1_D_ and not Rac1_T_ is extracted from the membrane, so that the Rac1 cycle is unidirectional (there is no branch that brings Rac1_T_ back to the cytoplasm, which would short-circuit the cycle). It has been proposed that tyrosine phosphorylation of membrane-bound Rac1 may facilitate its binding to RhoGDI [41], ensuring the unidirectionality of its GTPase cycle, similar to what has been shown for Ras [42]. Indeed, the release of membrane-bound Rac1_D_ into the cytoplasm has been shown to be favored by an order of magnitude over Rac1_T_ in both GDI-dependent and GDI-independent extraction mechanisms [43,44]. Therefore, it is justified to neglect membrane-bound Rac1_T_ in the model, even though some minor “leakage” into the cytoplasm might occur. Next, we assume that Rac1-specific GAP is recruited to the cell membrane by Rac1_T_, similar to what has been proposed for Ras1 and Cdc42 [45,46]. Consistent with this, the localization of *D. discoideum* RhoGAPs Dd5P4 and CARMIL-GAP is largely cytosolic, and they are only transiently recruited to the membrane by interacting with activated Rac1 [47,48], in agreement with data available for mammalian RhoGAPs [49,50].

Cycle (ii) describes the interaction of Rac1_T_ with its effector DGAP1. Our model assumes that DGAP1, in contrast to RhoGAP, first binds to the membrane and then interacts with Rac1_T_ (Fig 3A, right-hand side). This assumption is supported by our previous finding that DGAP1 localizes to the cell membrane in cells with actin cytoskeleton disrupted by latrunculin A, in which Rac1 is displaced from the membrane, suggesting a Rac1-independent mechanism for DGAP1 membrane recruitment [14]. Two degradation pathways for the Rac1_T_-DGAP1 complex have been proposed. The first describes the release of both components into the cytosol in conjunction with the deactivation of Rac1. Several reports have shown that the DGAP1-related protein IQGAP1 negatively regulates Rac1 activity mediated by RacGAP1 [51]. In addition to the degradation of the Rac1_T_-DGAP1 complex, which leads to the release of both components into the cytosol, we have introduced another degradation pathway in which Rac1_T_ remains bound to the membrane. Biologically, this corresponds to the possibility that a single Rac1 molecule sequentially binds multiple effectors before being released into the cytoplasm.

Based on the described interaction scheme, we have created a mathematical model that includes a series of reactions obeying the law of mass action. The results of computer simulations suggest that pattern formation is based on the accumulation of Rac1 at the membrane and its dissociation from the membrane stimulated by the interaction with GAP or DGAP1. The model reproduces all observed types of Rac1 dynamics: rotations, oscillations and stably polarized states, while preserving the phase relationships between the Rac1 and DGAP1 density profiles. The model is one-dimensional, where the position *x* represents a location at the cell edge and lies within the interval (0, *L*), where the symbol *L* represents the cell perimeter (Fig 3A). This is an appropriate choice for cell geometry because active Rac1, which drives cell motility, accumulates predominantly at the cell edge. For simplicity, we have also chosen the one-dimensional geometry for the cytoplasm.

In our mean-field approach, we describe the distributions of proteins along the cell perimeter by linear densities, where the distributions for the membrane-bound active Rac1, DGAP1, Rac1_T_-GAP complex and Rac1_T_-DGAP1 complex are labelled *R*_*T*_, *D*_*m*_, *G*_*c*_ and *D*_*c*_, respectively. We also describe the cytoplasmic distributions of inactive Rac1, GAP and DGAP1 by linear densities, denoted by *R*_*D*_, *G* and *D*, respectively. Since the positions *x* = 0 and *x* = *L* correspond to the same physical point on the cell membrane, we impose a periodic boundary condition on all densities and their first spatial derivatives, including those at the membrane and in the cytoplasm.

The cycle (i) is described by the following reaction-diffusion equations:

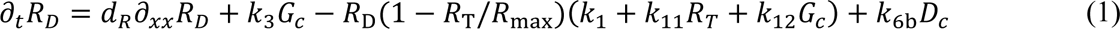

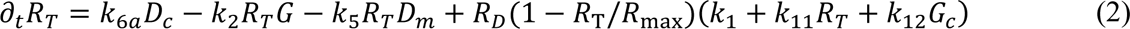

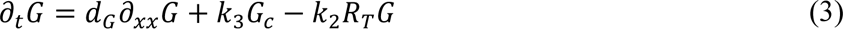

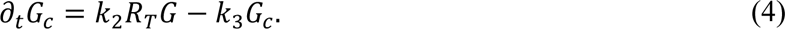

Here the reactions comprise several steps, with the symbol *k* indicating the reaction rate constant: binding of cytoplasmic Rac1_D_ to the membrane and its activation by GEF, *k*_1_, binding and activation stimulated by membrane-bound Rac1_T_, *k*_11_, and by the Rac1_T_-GAP complex, *k*_12_; binding of cytoplasmic GAP to Rac1_T_, leading to the formation of the Rac1_T_-GAP complex, *k*_2_; deactivation of Rac1 within the Rac1_T_-GAP complex and subsequent release of both proteins into the cytoplasm, *k*_3_. The model also includes the cytoplasmic diffusion of Rac1 and GAP; the corresponding diffusion constants are denoted by *d*_*R*_ and *d*_*G*_, while diffusion along the membrane is neglected. The term (1 − *R*_T_/*R*_max_) in Eq. (1b) ensures that the membrane density of Rac1 cannot exceed a saturation value, *R*_max_.

The cycle (ii) is described by the following reaction-diffusion equations:

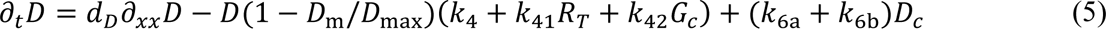

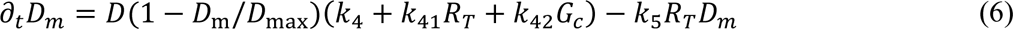

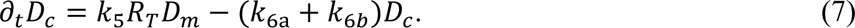

The reactions of this cycle include: attachment of cytoplasmic DGAP1 to the membrane, *k*_4_, attachment stimulated by Rac1_T_, *k*_41_, and by the Rac1_T_-GAP complex, *k*_42_; formation of the Rac1_T_-DGAP1 complex from membrane-bound DGAP1 and Rac1_T_, *k*_5_; degradation of the Rac1_T_-DGAP1 complex via two different pathways: release of DGAP1 into the cytoplasm, leaving Rac1_T_ at the membrane, *k*_6a_, or release of both proteins into the cytoplasm, *k*_6b_. We include the diffusion of DGAP1 molecules in the cytoplasm with a diffusion constant *d*_*D*_, while we neglect their diffusion at the membrane. The term (1 − *D*_m_⁄*D*_max_) in Eq. (3b) prevents excessive accumulation of DGAP1 molecules at the membrane with a saturation value of density, *D*_max_. Equations (1)-(7) together represent the joint description of the dynamics of Rac1, GAP and DGAP1.

### Patterns generated by the model

To investigate the behavior of our system, we solved the model numerically for biologically relevant parameters (S2 Appendix Table 1). Our results show that after a transient phase, the system converges to either a dynamic (traveling or standing waves) or a stationary regime (Figs. 3B-3D). In the case of traveling waves, all membrane-bound protein distributions are characterized by distinct peaks moving at the same constant speed (cf. line profiles for 3 time points in Fig 3B). The profiles of Rac1_T_ and the Rac1_T_-GAP complex form similar patterns, with Rac1T-GAP lagging slightly behind. In contrast, Rac1_T_ and DGAP1 are almost anti-correlated, with the minimum of Rac1_T_ near the maximum of DGAP1 and vice versa. Since the traveling waves occur along the closed loop, they correspond to the rotational movement of protein distributions along the cell membrane. For standing wave patterns, Rac1_T_ and DGAP1 are again anti-correlated, while the relationship between Rac1_T_ and Rac1_T_-GAP is more complex than for traveling wave patterns (Fig 3C). Typically, the appearance of a Rac1_T_ density peak is accompanied by a local minimum flanked by two maxima of the Rac1_T_-GAP complex distribution. The subsequent disappearance of the Rac1_T_ peak is accompanied by the formation of a new peak at the position shifted by *L*/2, which corresponds to the opposite side of the cell. Stationary patterns are characterized by sharply separated areas with different protein densities. For Rac1_T_ and the Rac1_T_-GAP complex, there is a large difference in density between the two domains, whereas the difference is smaller for DGAP1 (Fig 3D). Remarkably, the correlation between the distributions of Rac1_T_ and DGAP1 can be either positive or negative, depending on the total number of DGAP1 molecules. In both cases, Rac1_T_ and Rac1_T_-GAP are co-localized.

To systematically investigate the parameter space, we performed a linear stability analysis of our model. We used Fourier modes to describe the spatial perturbation of the steady state and examined the stability of each mode individually. By varying the total number of Rac1 and DGAP1 molecules (*N*_*R*_ and *N*_*D*_), we discovered two types of instabilities and the corresponding regions in parameter space: Hopf instability (oscillatory) and stationary instability (Fig 3E *left*). In the Hopf region, only the first Fourier mode is unstable, indicating that the model can generate oscillatory monopoles under these conditions. Conversely, in the stationary region, we observed areas where either only the first Fourier mode was unstable or all Fourier modes were unstable. This overall instability of all Fourier modes can drive the system into a multipolar state where the number of poles is directly influenced by the initial protein distribution. We then repeated the systematic exploration of the parameter space, but this time using large-scale computer simulations instead of linear stability analysis. The patterns resulting from these simulations were then plotted based on the total number of Rac1 and DGAP1 molecules (Fig 3E *right*). The map derived from the simulation and the map from the linear stability analysis showed significant similarity, except for the high *N*_*R*_ and high *N*_*D*_region, where the linear stability analysis predicted the formation of periodic patterns but the simulations resulted in stationary patterns.

### Spatio-temporal dynamics of Rac1 activity: a comparison between experimental and modeled results

To compare the patterns obtained experimentally and theoretically, we juxtaposed kymographs of the measured Rac1* signal in the membrane and the linear concentration of Rac1_T_ calculated by the model for three typical cases (Fig 4). In addition to kymographs, we also generated corresponding autocorrelograms to facilitate pattern recognition (Fig 4A**–**B, S3 Fig A). In the kymographs and autocorrelograms, the traveling waves of the Rac1_T_ distribution are represented as parallel stripes whose slope reflects the wave velocity (Fig 4A). Oscillations lead to regular patterns of patches, while stationary patterns appear as vertical stripes (Figs 4B and 4C). The model reproduces major characteristics of experimentally observed patterns of Rac1 activity. We found that with an appropriate choice of parameter values, the model reproduces the typically measured velocities and wavelengths of traveling waves, the periods of oscillation of monopolar standing waves, and the contrast between and relative widths of active and inactive domains in stationary monopolar patterns. In addition to reproducing the typically observed patterns, we have also successfully simulated less frequently occurring patterns, such as oscillating dipoles (S3 Fig A), rotating dipoles (S3 Fig B) and stationary dipoles (S3 Fig C). In order to make experimentally testable predictions, we investigated how the oscillation patterns depend on the cell perimeter *L*. Linear stability analysis predicts that higher Fourier modes become unstable with increasing *L* (S4 Fig A). We tested this prediction by measuring the perimeters of cells that exhibited monopolar and dipolar patterns. Our results showed that monopolar patterns occurred predominantly in small cells, while dipolar patterns occurred in larger cells (S4 Fig B).

**Fig 4.**
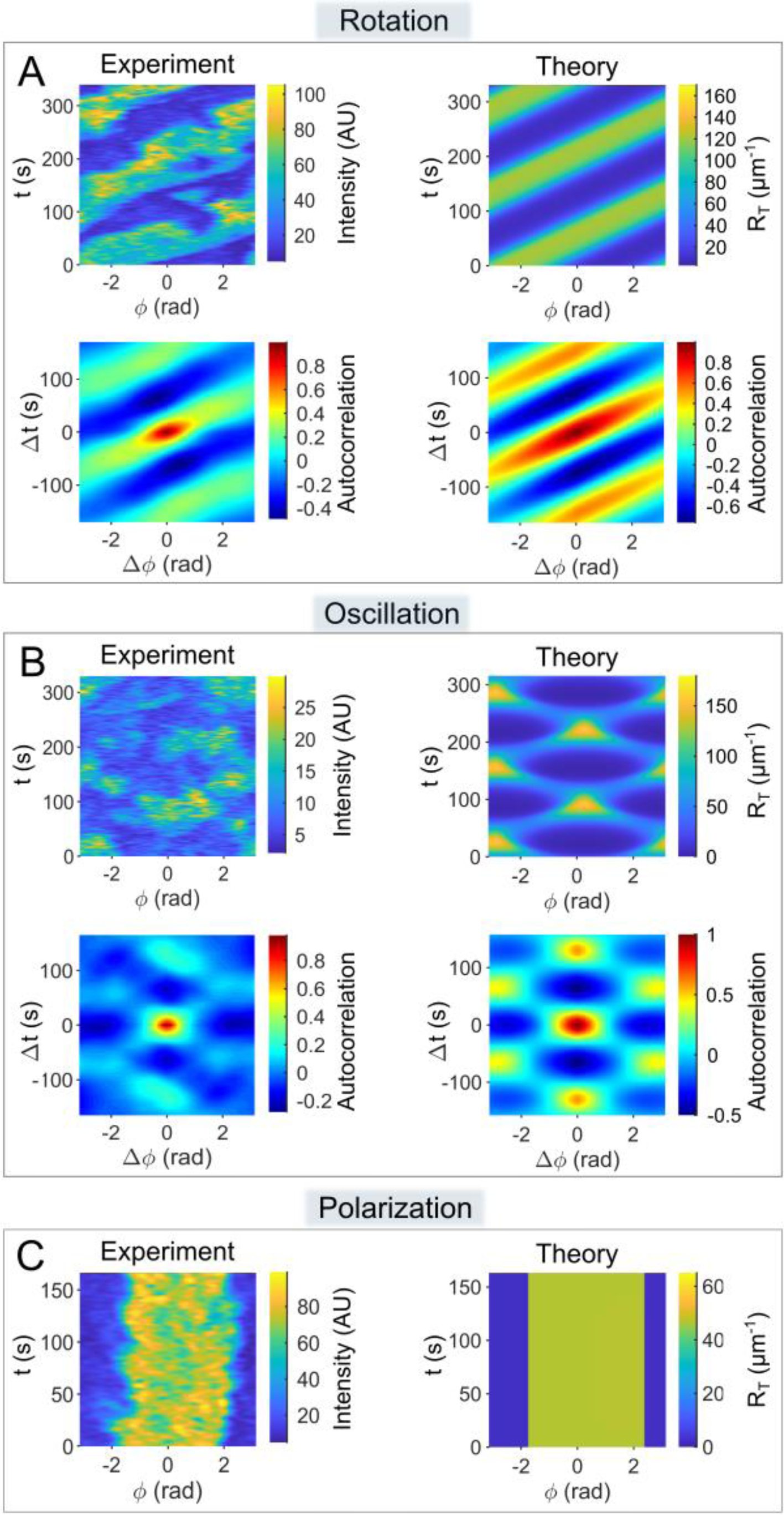
Comparison of the experimentally observed patterns of Rac1* with the patterns of Rac1_T_ derived from computer simulations. In panels A-B, the patterns are shown as kymographs (using the Matlab colormap parula) next to the corresponding autocorrelograms (Matlab colormap: jet). Only kymographs are shown in panel C. The experimental kymographs show the fluorescence intensity measured along the cell perimeter (angular position, *ϕ*) versus time (*t*). Equivalent maps of the calculated linear Rac1_T_ concentration, *R*_*T*_, are shown in the theoretical kymographs. Autocorrelograms represent the self-product of appropriately normalized kymographs that are shifted in both time (Δ*t*) and space (Δ*ϕ*). A precise definition can be found in the Material and methods section. The patterns displayed correspond to the dynamics types shown in Fig 1: (**A**) *Rotating monopole*, (**B**) *Oscillating monopole* and (**C**) *Stationary monopole*. The parameter values used in the simulations are listed in S2 Appendix Table 1.

In our experiments, we not only observed transitions between ordered and random patterns, but also transitions between different ordered patterns. For example, we observed a transition from oscillation to rotation, as well as a transition from clockwise to counterclockwise rotation (Fig 5). These results prompted us to investigate whether such transitions can also occur in simulations. Our numerical simulations show that oscillations can transition to rotations and that the duration of these transitions is on the time scale of one period (Fig 5A). In addition, our simulations also show transitions from clockwise to counterclockwise rotations on a similar time scale for different parameter values (Fig 5B).

**Fig 5.**
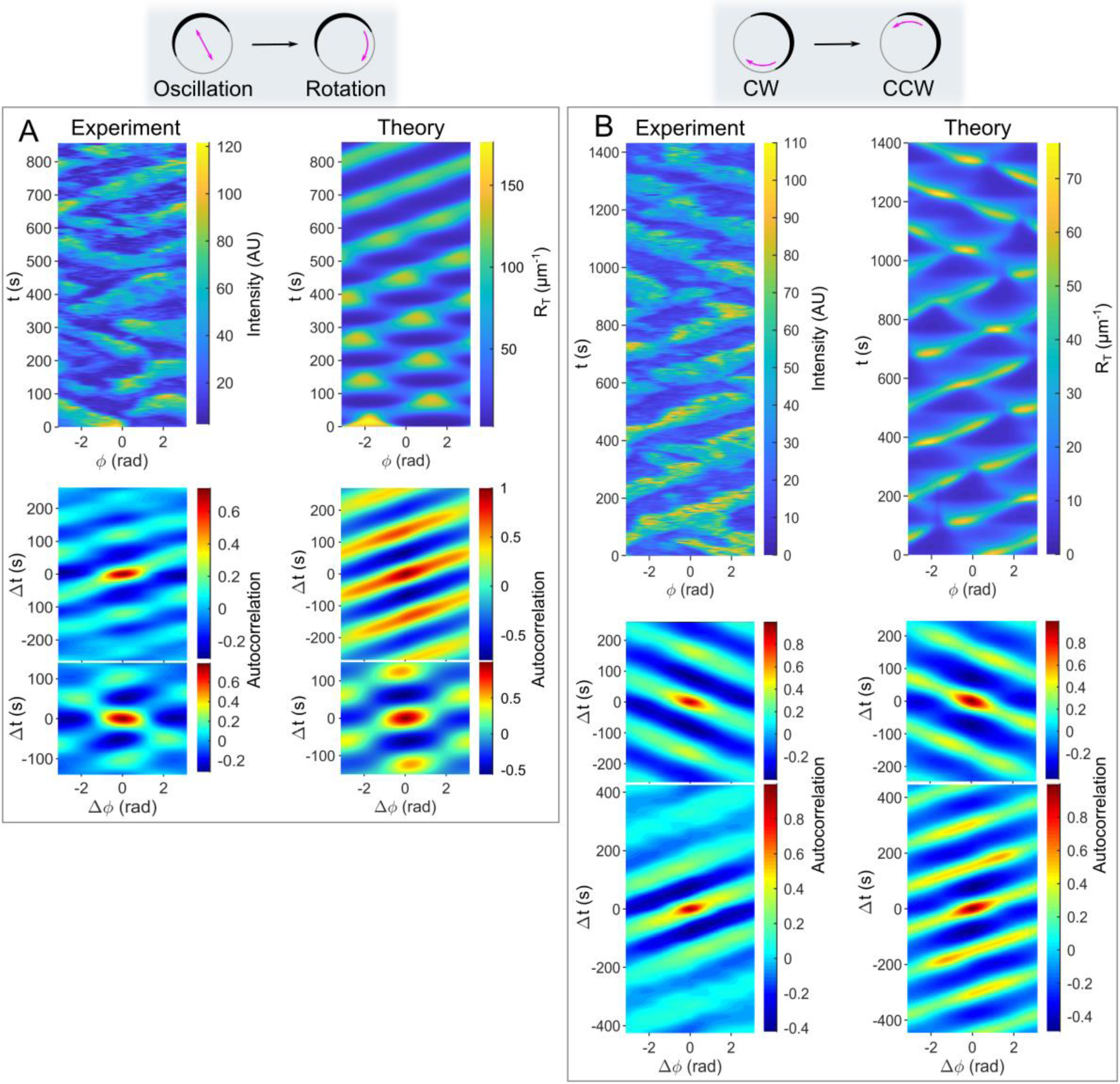
Transitions between different dynamic regimes. (**A**) An experimentally observed transition between the oscillation and the rotation of a single domain/monopole (*left*), and the corresponding theoretical simulation (*right*). (**B**) An experimentally observed transition between the clockwise and counterclockwise rotation of a single domain/monopole (*left*) and the corresponding theoretical simulation (*right*). In this scenario, some parameter values were significantly adjusted from their original estimates to bring the period of the traveling waves in line with the observed experimental data (see S2 Appendix–Table 1). Kymographs and autocorrelograms are defined as described in Fig 4.

### Phase relationships between Rac1T and its effector DGAP1

Since both DPAKa(GBD)-DYFP and DGAP1-mRFP interact with active Rac1, it’s counterintuitive to observe their fluorescence signals on opposite sides of the cell (Fig 2). To contrast the experimentally observed negative correlation with our theoretical model, we compared kymographs of Rac1* and DGAP1* from microscopy with our theoretical predictions. Both the experimental and theoretical data showed primarily anti-correlated distributions between Rac1 and DGAP1 in oscillating patterns (Fig 6A). To further investigate the relationship between Rac1* and DGAP1* during rotation, we analyzed time series corresponding to specific locations at the cell edge. However, analyzing fluorescence from a single spatial point proved impractical due to excessive noise in the data series. Therefore, we decided to use averaged time series obtained from the data of all spatial points in a single experiment. First, we had to align all time series corresponding to the different locations on the circumference to account for their mutual phase shifts. For this matching, we used principal component analysis (see Appendix 3). After obtaining the characteristic temporal fluorescence intensity profiles for Rac1* and DGAP1*, we compared them to the model simulation (Fig 6B). The calculated profiles agreed remarkably well with the experimental data in terms of periodicity, asymmetry, relative intensities and the phase relationship between the two waves. Using the same experimental data sets, we created a phase portrait showing the counterclockwise evolution along a crescent-shaped trajectory (Fig 6C, *left*). The phase portrait derived from our model simulations mirrors that of the experimental data. To capture the correct counterclockwise evolution in the phase portrait, we included the cooperative binding of DGAP1 to the membrane, which is described in the model equations by terms with constants *k*_41_and *k*_42_. Without these terms, the system always evolved in a clockwise direction, regardless of the choice of parameter values.

**Fig 6.**
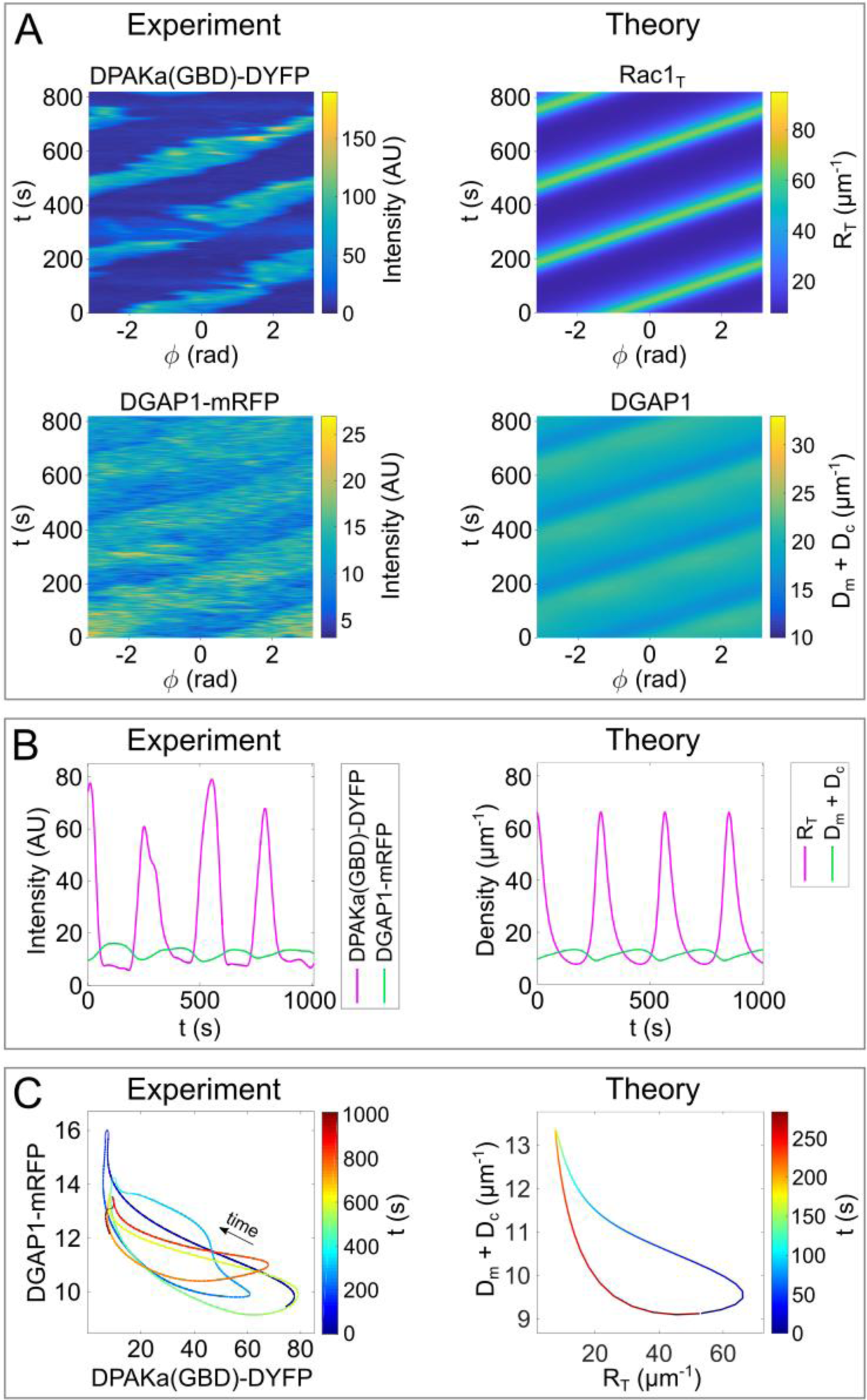
Relationship between the travelling waves of Rac1 and DGAP1. (**A**) Experimentally observed rotational patterns of Rac1* and DGAP1* next to the corresponding patterns of Rac1_T_ and DGAP1 derived from computer simulations. (**B**) *Left* The average fluorescence intensities of Rac1* and DGAP1*, as obtained by principal component analysis of travelling waves, plotted as functions of time. *Right* The time series of Rac1_T_ and DGAP1 densities at a spatial point as derived from computer simulations. (**C**) Phase portraits created using the data points from (B). The phase portrait derived from the simulated data represents a rotation period (limit cycle). In both experiment and theory, the system moves counterclockwise along a crescent-shaped trajectory. The time course is represented by color coding. The parameters correspond to the values given in S2 Appendix–Table 1.

## DISCUSSION

Our study reveals intricate spatiotemporal dynamics of Rac1 activity in *D. discoideum*, comprising periodic and irregular patterns as well as transitions between them (Figs 1 and 5). The observed oscillation patterns exhibit periodicities typically between 2 and 5 minutes. Comparable intracellular oscillations of small Rho-family GTPases have been observed in a whole spectrum of eukaryotic organisms, from yeast to humans [52]. In budding yeast, clusters of active Cdc42 oscillate with a characteristic period of 4-5 minutes [53]. In fission yeast cells, which grow bipolar, active Cdc42 oscillates between the polarized cell tips with an average period of 5 minutes [54]. In human NK cells, Cdc42 activity oscillates with a periodicity of 5 to 6 minutes [55]. In developing *Xenopus* embryos, active Rho forms waves, while in reconstituted *Xenopus* cortices it organizes in oscillating patches with characteristic periodicities of 1 to 3 minutes [56,57]. Similarly, in plants, the active GTPase Rop1 undergoes oscillatory accumulation and spreading at the tips of growing pollen tubes with a periodicity between 1 and 3 minutes [58]. These consistent periodicities in different organisms suggest that common regulatory mechanisms control the dynamics of Rho GTPases.

Early studies found that the shape changes during locomotion of *D. discoideum* cells are not random [59,60]. Quantitative analyses revealed elongations, rotations and oscillations in the shape of both vegetative and starved *D. discoideum* cells and showed a correlation with the dynamics of F-actin [61]. The periodicity of the observed rotations and oscillations was between 2.5 and 3.5 minutes, which is consistent with our results. Each type of recurrent pattern lasted 10 to 20 minutes, and the patterns occasionally transitioned spontaneously. However, no bipolar oscillations or rotations of F-actin were detected. Our work and others link Rac activation to actin-driven protrusions in *D. discoideum* and other cells [14,22,35,62]. Therefore, it is highly plausible that the dynamic phenomena observed by Maeda and colleagues are related to those described in this study. However, while transitions between different periodic F-actin patterns were hypothesized to occur stochastically, here we show that transitions in active Rac1 patterns can occur within the deterministic framework of our model (Fig 5).

In mutant *D. discoideum* cells lacking enzymes that regulate the interconversion of phosphoinositides (PtdIns), such as PI3K and PTEN, the periodic changes in cell shape were significantly reduced [61]. Subsequently, oscillations in the PtdIns signaling system were observed using fluorescently labelled PIP3 and PTEN in cells immobilized with latrunculin A and treated with caffeine [63]. These studies revealed that the observed oscillations and rotations of membrane domains differentially enriched with PTEN or PIP3 exhibit the properties of a relaxation oscillator [63,64]. Links between PtdIns and Rho-GTPase signaling and their oscillations have been suggested in various cellular systems. For example, in neutrophils, PIP3 and active Rac promote actin polymerization, which in turn drives PIP3 and Rac activity through reciprocal positive feedback mechanisms [65–68]. In natural killer cells that form immunological synapses with target cells, the p85a subunit of phosphoinositide 3-kinase (PI3K) was required to maintain oscillations in Cdc42 activity [55]. Given the striking similarity between the oscillations of PIP3 and Rac1 in *D. discoideum*, the question arises whether the established pattern of the former might serve as a template for the spatiotemporal distribution of the latter [69]. Direct experimental testing of this proposition in *D. discoideum* is challenging, as the PIP3 waves were observed in immobilized cells treated with latrunculin A and caffeine, a treatment that abolishes the F-actin-dependent cortical localization of Rac1 [14,35]. However, our theoretical studies suggest that the autonomous signaling system, which involves the activation and inactivation of Rac1 and its switching between the membrane and the cytosol, is capable of generating different oscillatory patterns independently of other signaling systems.

Rho GTPases play a crucial role in coordinating the timing, location and nature of supramolecular actin assemblies in motile cells [36,70]. Interestingly, a single Rho GTPase can regulate different cytoskeletal activities by selectively interacting with different downstream effectors [71]. It has been shown that RacA in mammals and the related Rac1 in *D. discoideum* control the protrusion of lamellipodia and pseudopodia [14,22,35,72], which is facilitated by Rac1-dependent activation of actin polymerases [2,6,7,73]. However, Rac1 also regulates non-protrusive and contractile areas of the actin cortex, albeit apparently to a lesser extent [13,74–76]. Our study contrasts the dynamics of a Rac1 effector, DGAP1, involved in actin filament entanglement in the posterior and lateral cortex, with the cortical distribution of Rac1*. We confirmed that cortical regions enriched in DGAP1 mostly do not overlap with those enriched in Rac1* [13]. Surprisingly, however, we also observed rare domains enriched in both proteins in polarized cells (Fig 2B), a feature supported by our theoretical model (Fig 3).

To investigate the observed dynamics from a theoretical perspective, we constructed a reaction-diffusion model based on the generally known interactions between Rac1, its associated regulators from the GAP, GEF and GDI families, and its effector DGAP1. To keep the model manageable, we only explicitly considered the distributions and mutual interactions between Rac1, GAP and DGAP1 (Fig 3A), while the activities of the other regulators were indirectly included in the interaction cycle of Rac1 (S1 Appendix). The model contains nonlinear terms describing feedback mechanisms that facilitate the recruitment of Rac1 and DGAP1 to the membrane. It successfully replicates the spatiotemporal distributions of active Rac1 and DGAP1 in rotational, oscillatory and polar states, transitions between the observed dynamic regimes, and the relative abundances and phase relationships between active Rac1 and DGAP1 on the membrane. Therefore, our computational model provides an explanation for the experimentally observed negative correlation between the concentration peaks of active Rac1 and its effector DGAP1 [13,14]. Remarkably, the occurrence of periodic Rac1 patterns in our simulations was independent of the presence of DGAP1. The model also predicts a tight coupling between the dynamics of active Rac1 and GAP, a hypothesis that could be tested by future experimental research on RacGAP candidates in *D. discoideum*.

In our model, we explicitly considered the interaction of Rac1 with only one of its many effectors [7]. However, the model accounts for positive cooperativity in the binding of Rac1 to the plasma membrane in conjunction with its activation. The underlying mechanism likely involves the interaction of Rac1 with other effectors that promote actin polymerization [40]. It is known that some effectors of Rho GTPases can directly modulate their activity within multi-protein complexes [24]. In general, autocatalytic, GEF-mediated and effector-assisted activation is a universal feature of Rho-GTPase regulation in a variety of eukaryotic cells [28,38]. RacGEF1 has also been shown to mediate a positive feedback loop between F-actin and Rac activity in *D. discoideum*, although the details of the underlying mechanism are still unclear [40].

Early models that attempted to explain the symmetry breaking in the distribution of Rho GTPases were inspired by the polarized clustering of Cdc42 in *S. cerevisiae* [27,77]. In these seminal studies, it was recognized that autocatalytic GEF-mediated activation is crucial for the polarization of Cdc42 [26]. A little later, active Cdc42 was found to oscillate between transient foci in *S. cerevisiae* and between incipient poles in *S. pombe* [53,54]. It is often assumed that oscillatory behavior in biological systems requires the presence of a delayed negative feedback loop [78]. Therefore, the oscillations of Cdc42 between the two poles in fission yeast cells were modeled under the assumption that Cdc42 triggers its own elimination through a generic delayed negative feedback mechanism [54]. In budding yeast, negative feedback mechanisms have been proposed to act through Cdc42-mediated activation of a Cdc42-driven GAP or through inhibitory phosphorylation of the Cdc42-specific GEF Cdc24 by the Cdc42 effector Pak1 [53,79,80]. A similar role for Cdc42-specific GEFs and GAPs in the regulation of oscillations by negative feedback was investigated in *S. pombe* [45,81].

However, recent studies have shown that in yeast, the stability of multipolar distributions of active Rho GTPases in mass-conserving reaction-diffusion models does not require the explicit introduction of nonlinear negative feedback [82,83]. Instead, the coexistence of multiple GTPase clusters is determined by the extent to which the maximum concentration of a limiting protein species approaches the “saturation point” determined by model parameters [82,84]. Therefore, it has been suggested that protein concentrations, reaction rate constants and cell size, rather than specific feedback mechanisms, determine whether unipolar or multipolar outcomes are achieved [82,83]. Furthermore, indirect pathways for the conversion of Rho-GTP to Rho-GDP, likely involving RhoGAPs, have been hypothesized to be critical for controlling local concentrations [84,85]. Our results suggest that a similar principle may control the occurrence and transformation of oscillations and polarized distributions of Rac1-GTP in *D. discoideum*.

In summary, our experimental results support the idea that the activities of small Rho GTPases are prone to autonomous, constitutive spatiotemporal oscillations in eukaryotic cells. Our theoretical model for Rac1 dynamics in *D. discoideum* cells contributes to the understanding of Rac1 regulation and membrane localization. It supports the idea that regulation of Rho-GTPase signaling activities by GEFs and GAPs can lead to oscillatory dynamics [86]. Future research should investigate the impact of the integrity of the PI-centered signaling network on Rac1 oscillations as well as the contribution of effectors beyond DGAP1 to Rac1 dynamics. Furthermore, our current model does not yet take into account the mechanical forces caused by actin polymerization and membrane deformation [25,87–89]. The functional significance of Rho-GTPase oscillations in general remains a controversial topic. Some propose that they represent an exploratory behavior typical of self-organizing biological systems [54,90], while others suggest that the oscillations may be a by-product of the topology of the interaction network [52,91]. In *D. discoideum* cells, Rac1 oscillations could facilitate the rapid adaptation of cytoskeletal activities to environmental changes. Future studies should investigate the effects of stochasticity on the extent of the different dynamic regimes and the transitions between them. Considering that regulatory and signaling mechanisms involving small Rho GTPases are largely conserved in eukaryotes [7,8,92], we hypothesize that our model is potentially applicable to Rac polarization and oscillations in other systems as well.

## Materials and methods

### Expression vectors and *D. discoideum* cell lines

*Dictyostelium discoideum* cells were cultivated in polystyrene culture dishes at 22 °C in axenic HL5 medium (Formedium), with the addition of 18 g/L maltose, 50 μg/ml ampicillin and 40 μg/ml streptomycin. We used AX2 cells stably expressing the DPAKa(GBD)-DYFP probe constructed as previously described [14]. For the co-expression of DPAKa(GBD)-DYFP and mRFP-DGAP1, AX2 cells stably expressing mRFP-DGAP1 were transfected with the pDEX-DPAKa(GBD)-DYFP vector [14]. Cell transfection by electroporation and clonal selection was performed as described [93].

### Confocal laser scanning fluorescence microscopy

Fluorescence microscopy was performed using a Leica TCS SP8 X microscope equipped with a supercontinuum excitation laser (Leica Microsystems). The excitation wavelengths and detection ranges used for imaging were: (1) 511 nm and 520-565 nm for DYFP, and (2) 575 nm and 585-630 nm for mRFP. The hybrid HyD detectors were operated in gated mode to suppress the parasite reflection from the coverslip surface, and the delay time between excitation and detection was set to 0.3 ns. Only an approximately 1 μm thick section of typically 5-10 μm thick cells was imaged, corresponding to the depth of field of the imaging setup. In total, we observed 85 randomly selected cells expressing DPAKa(GBD)-DYFP and 93 cells co-expressing DPAKa(GBD)-DYFP and DGAP1-mRFP. After selecting a cell that expressed fluorescent probe(s) at moderate levels, it was recorded for at least 10 minutes, and if no obvious patterns were visible, the cell was categorized as having no spatially structured dynamics. Otherwise, the observed pattern was further analyzed. Of the 85 monitored cells expressing DPAKa(GBD)-DYFP, the dynamics in 59 cells were classified as random. Of the 93 monitored cells co-expressing DPAKa(GBD)-DYFP and DGAP1-mRFP, the dynamics were classified as random in 59 cells. In the following analyses of DPAKa(GBD)-DYFP dynamics, the results of both cell types were combined.

### Image processing

QuimP, a set of plugins for ImageJ (NIH), was used to analyze the intensities of the fluorescent probes on the cell cortex [94]. To remove excess noise from the data, we loaded the output intensities into Matlab (MathWorks) and processed the data with the smoothing spline function `csaps`. The resulting variable *I*(*ϕ*, *t*) was then color-coded and plotted against the angle *ϕ*, which represents the position on the cell membrane, and time *t*. To facilitate the detection of repetitive patterns reflecting protein dynamics, we calculated the autocorrelation function of *I*(*ϕ*, *t*). We define the autocorrelation function of fluorescence intensity *I* as

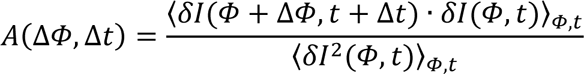

where *δI*(*ϕ*, *t*) = *I*(*ϕ*, *t*) − ⟨*I*(*ϕ*, *t*)⟩_*ϕ*_, and ⟨ ⟩_*ϕ*,*t*_ denotes the average over the angle and time.

### Numerical methods

For the numerical calculations, the spatial domain was discretized into 100 uniformly distributed points given by the vector *x* = (*x*_1_, …, *x*_100_), where each component has a value *x*_*i*_ = (*i* − 1) · ℎ, with index *i* = 1, …, 100 and distance between adjacent spatial points ℎ = *L*/100. In this case, the spatial derivatives have an approximate value:

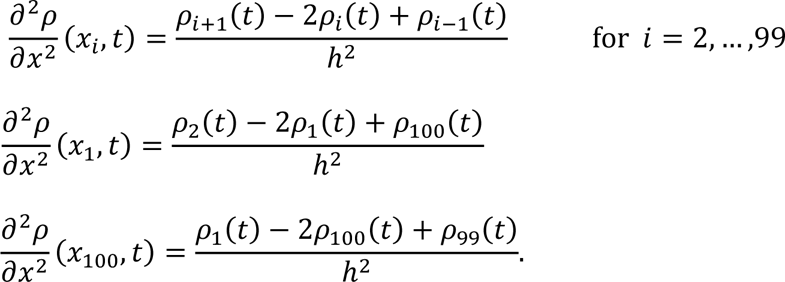

Here, the vector *ρ* = (*R*_*D*_, *R*_*T*_, *G*, *G*_*c*_, *D*, *D*_*m*_, *D*_*c*_)^*T*^ represents the densities of the protein species. The periodic boundary condition used in this study is included in the equations for the boundaries at *i* = 1 and *i* = 100. Consequently, each partial differential equation was replaced by a system of 100 coupled ordinary differential equations. We solve such a system using Matlab solver `ode15s`, a variable-step, variable-order method based on backward difference formulas.

### Linear stability analysis

To investigate the nature of the solutions in larger parts of the parameter space, we performed a linear stability analysis. The original reaction-diffusion system, given by Eqs. (1)-(7), can be expressed in a compact form

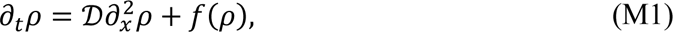

where *f*: ℝ^7^ → ℝ^7^ is a nonlinear function given by:

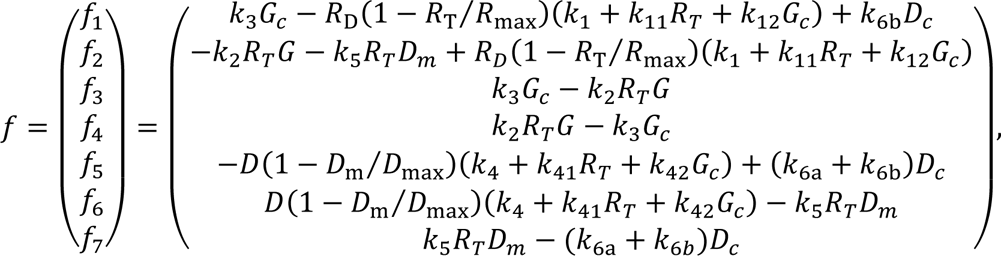

and *D* is 7 × 7 diagonal matrix containing diffusion constants,

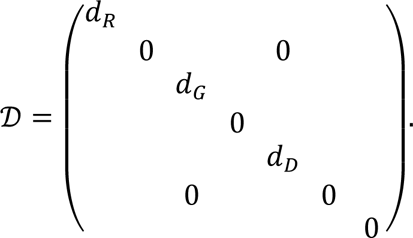

We have first determined a fixed point of the system, which is referred to as *ρ*^∗^. For the spatially homogeneous fixed point, Eq. (M1) reduces to *f*(*ρ*^∗^) = 0. These seven equations are linearly dependent, with four of them being independent:

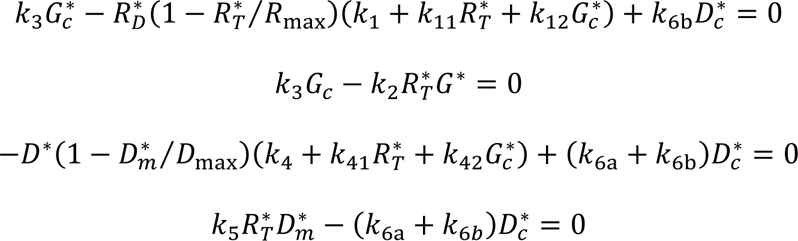

Furthermore, the total number of molecules of each protein remains constant, regardless of whether these molecules are in a complex or free:

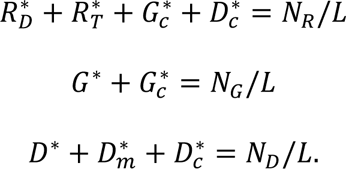

Here, the symbols *N*_*R*_, *N*_*G*_, and *N*_*D*_represent the total number of Rac1, GAP, and DGAP1 molecules, respectively. For a given set of parameter values, these equations have a unique solution, which we have calculated using Matlab.

To assess the stability of the steady state, we have considered the linearized system

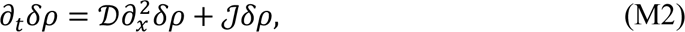

where *δρ* = *ρ* − *ρ*^∗^ represents the deviation from the steady state, and *J* denotes the Jacobian matrix of *f*, which is defined as

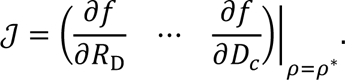

The linearized equations (M2) can be solved using the ansatz *δρ*_*q*_ ∝ *e*^*σ*^_*q*_^*t*+*iqx*^. The stability of the steady state depends on the growth rate *σ*_*q*_, whereby a positive value of the real part indicates an unstable state. The symbol *q* stands for the wavenumber, whose possible values are limited by the boundary conditions. For periodic boundary conditions, the wavenumbers are expressed as *q* = 2*nπ*/*L*, where *n* ∈ {0, ±1, ±2, … }. By inserting the ansatz into the linearized system, a connection between the values of *σ*_*q*_ and *q* is established:

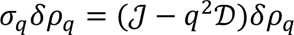

For a given wavenumber, the values of *σ*_*q*_ are therefore the eigenvalues of the matrix ℳ_*q*_ = *J* − *q*^2^*D*. If the real part, Re*σ*_*q*_, becomes positive when a control parameter is changed and the imaginary part, Im*σ*_*q*_, is not equal to zero, an oscillatory or Hopf bifurcation occurs. If Im*σ*_*q*_ = 0, however, the instability is described as stationary.

## Author Contributions

**Conceptualization**: Marko Šoštar, Nenad Pavin, Igor Weber.

**Formal analysis**: Marko Šoštar, Nenad Pavin, Igor Weber.

**Funding acquisition**: Igor Weber, Nenad Pavin.

**Investigation**: Marko Šoštar.

**Methodology**: Marko Šoštar, Maja Marinović, Vedrana Filić, Nenad Pavin, Igor Weber.

**Project administration**: Nenad Pavin, Igor Weber.

**Software**: Marko Šoštar.

**Supervision**: Nenad Pavin, Igor Weber.

**Visualization**: Marko Šoštar, Nenad Pavin.

**Writing – original draft**: Marko Šoštar, Nenad Pavin, Igor Weber.

**Writing – review & editing**: Marko Šoštar, Maja Marinović, Vedrana Filić, Nenad Pavin, Igor Weber.

## Supporting information

**S1 Movie**. **Representative rotating monopole.** A Rac1*-enriched domain travels along the cell membrane (corresponds to Fig 1A).

**S2 Movie**. **Representative rotating dipole.** Two distinct Rac1*-enriched domains travel along the cell membrane (corresponds to S1 Fig A).

**S3 Movie**. **Representative oscillating monopole.** A Rac1*-enriched domain periodically relocates from one side of the cell to the opposite side (corresponds to Fig 1B).

**S4 Movie**. **Representative oscillating dipole**. Two distinct Rac1*-enriched domains periodically undergo 90-degree orientation shifts (corresponds to S1 Fig B).

**S5 Movie**. **Representative stationary monopole.** A Rac1*-enriched domain remains on one side of a migrating cell (corresponds to Fig 1C).

**S6 Movie**. **Representative stationary dipole.** Two distinct Rac1*-enriched domains located on opposite sides of the cell maintain their positions (corresponds to S1 Fig C).

**S7 Movie**. **Representative rotating monopoles.** Rac1* and DGAP1-enriched domains, located on opposite sides of the cell, travel along the cell membrane (corresponds to Fig 2A).

**S8 Movie**. **Representative oscillating dipoles.** Two distinct Rac1*-enriched domains and oppositely localized DGAP1-enriched domains periodically shift their orientation by 90 degrees (corresponds to S2 Fig).

**S9 Movie**. **Representative stationary monopoles out of phase**. Rac1*-enriched domain and oppositely localized DGAP1-enriched domain maintain their positions (corresponds to Fig 2B).

**S10 Movie**. **Representative stationary monopoles in phase.** Rac1*-enriched and DGAP1-enriched domains are co-localized, maintaining their position on the cell membrane (corresponds to Fig 2C).

**S1 Figure**. **Representative bipolar patterns of cortical domains enriched in Rac1*.**

**S2 Figure**. **A representative oscillating dipole in a double-labeled cell.**

**S3 Figure**. **Comparison of the experimentally observed patterns of Rac1* with the patterns of Rac1_T_ derived from computer simulations.**

**S4 Figure**. **Effect of circumference size on pattern formation.**

**S1 Appendix**. **Justification of the most important reaction mechanisms contained in the model.**

**S2 Appendix**. **Estimation of the values of parameters used in the simulations.**

**S3 Appendix**. **Principal component analysis.**

**S1 Supporting Information. MATLAB codes used to generate data shown in Figures 3, 4, 5, 6, S3 and S4.**

**S2 Supporting Information. Data obtained from the QUIMP analysis of the cell outline fluorescence intensity used as the input for MATLAB routines to generate Figures 4, 5, 6, S3 and S4.**

**S3 Supporting Information. Confocal microscopy series used to generate Figure 1A.**

**S4 Supporting Information. Confocal microscopy series used to generate Figure 1B.**

**S5 Supporting Information. Confocal microscopy series used to generate Figure 1C.**

**S6 Supporting Information. Confocal microscopy series used to generate Figure S1A.**

**S7 Supporting Information. Confocal microscopy series used to generate Figure S1B.**

**S8 Supporting Information. Confocal microscopy series used to generate Figure S1C.**

**S9 Supporting Information. Confocal microscopy series used to generate Figure 2A.**

**S10 Supporting Information. Confocal microscopy series used to generate Figure 2B, mRFP channel.**

**S11 Supporting Information. Confocal microscopy series used to generate Figure 2B, YFP channel.**

**S12 Supporting Information. Confocal microscopy series used to generate Figure 2C.**

**S13 Supporting Information. Caption to Supporting Information files 1-12.**

## References

1. Ishikawa-Ankerhold HC, Müller-Taubenberger A. Actin assembly states in Dictyostelium discoideum at different stages of development and during cellular stress. Int J Dev Biol. 2019;63(8-9–10):417–27.

2. Schaks M, Giannone G, Rottner K. Actin dynamics in cell migration. Essays Biochem. 2019 Oct 31;63(5):483–95.

3. Mosaddeghzadeh N, Ahmadian MR. The RHO Family GTPases: Mechanisms of Regulation and Signaling. Cells. 2021 Jul 20;10(7):1831.

4. Amano M, Nakayama M, Kaibuchi K. Rho-kinase/ROCK: A key regulator of the cytoskeleton and cell polarity. Cytoskelet Hoboken NJ. 2010 Sep;67(9):545–54.

5. Ding B, Yang S, Schaks M, Liu Y, Brown AJ, Rottner K, et al. Structures reveal a key mechanism of WAVE regulatory complex activation by Rac1 GTPase. Nat Commun. 2022 Sep 16;13(1):5444.

6. Kühn S, Geyer M. Formins as effector proteins of Rho GTPases. Small GTPases. 2014;5:e29513.

7. Filić V, Mijanović L, Putar D, Talajić A, Ćetković H, Weber I. Regulation of the Actin Cytoskeleton via Rho GTPase Signalling in Dictyostelium and Mammalian Cells: A Parallel Slalom. Cells. 2021 Jul;10(7):1592.

8. Forbes G, Schilde C, Lawal H, Kin K, Du Q, Chen ZH, et al. Interactome and evolutionary conservation of Dictyostelid small GTPases and their direct regulators. Small GTPases. 2022 Jan;13(1):239–54.

9. Rivero F, Xiong H. Rho Signaling in Dictyostelium discoideum. Int Rev Cell Mol Biol. 2016;322:61–181.

10. Litschko C, Linkner J, Brühmann S, Stradal TEB, Reinl T, Jänsch L, et al. Differential functions of WAVE regulatory complex subunits in the regulation of actin-driven processes. Eur J Cell Biol. 2017 Dec;96(8):715–27.

11. Mondal S, Burgute B, Rieger D, Müller R, Rivero F, Faix J, et al. Regulation of the Actin Cytoskeleton by an Interaction of IQGAP Related Protein GAPA with Filamin and Cortexillin I. PLOS ONE. 2010 Nov 10;5(11):e15440.

12. Faix J, Weber I, Mintert U, Köhler J, Lottspeich F, Marriott G. Recruitment of cortexillin into the cleavage furrow is controlled by Rac1 and IQGAP-related proteins. EMBO J. 2001 Jul 7;20(14):3705.

13. Filić V, Marinović M, Faix J, Weber I. The IQGAP-related protein DGAP1 mediates signaling to the actin cytoskeleton as an effector and a sequestrator of Rac1 GTPases. Cell Mol Life Sci CMLS. 2014 Aug;71(15):2775–85.

14. Filić V, Marinović M, Faix J, Weber I. A dual role for Rac1 GTPases in the regulation of cell motility. J Cell Sci. 2012 Jan 15;125(Pt 2):387–98.

15. Donnelly SK, Bravo-Cordero JJ, Hodgson L. Rho GTPase isoforms in cell motility: Don’t fret, we have FRET. Cell Adhes Migr. 2014 Oct 31;8(6):526–34.

16. Hodgson L, Pertz O, Hahn KM. Design and Optimization of Genetically Encoded Fluorescent Biosensors: GTPase Biosensors. In: Methods in Cell Biology [Internet]. Elsevier; 2008 [cited 2023 Feb 22]. p. 63–81. Available from: https://linkinghub.elsevier.com/retrieve/pii/S0091679X08850042

17. Martin K, Reimann A, Fritz RD, Ryu H, Jeon NL, Pertz O. Spatio-temporal co-ordination of RhoA, Rac1 and Cdc42 activation during prototypical edge protrusion and retraction dynamics. Sci Rep. 2016 Feb 25;6(1):21901.

18. Welch CM, Elliott H, Danuser G, Hahn KM. Imaging the coordination of multiple signalling activities in living cells. Nat Rev Mol Cell Biol. 2011 Nov;12(11):749–56.

19. Dagliyan O, Dokholyan NV, Hahn KM. Engineering proteins for allosteric control by light or ligands. Nat Protoc. 2019 Jun;14(6):1863.

20. Inoue T, Heo WD, Grimley JS, Wandless TJ, Meyer T. An inducible translocation strategy to rapidly activate and inhibit small GTPase signaling pathways. Nat METHODS. 2005;2(6):415–8.

21. Shcherbakova DM, Cox Cammer N, Huisman TM, Verkhusha VV, Hodgson L. Direct multiplex imaging and optogenetics of Rho GTPases enabled by near-infrared FRET. Nat Chem Biol. 2018 Jun 1;14(6):591–600.

22. Wu YI, Frey D, Lungu OI, Jaehrig A, Schlichting I, Kuhlman B, et al. A genetically encoded photoactivatable Rac controls the motility of living cells. Nature. 2009 Sep;461(7260):104–8.

23. Fritz RD, Pertz O. The dynamics of spatio-temporal Rho GTPase signaling: formation of signaling patterns. F1000Research. 2016 Apr 26;5:F1000 Faculty Rev-749.

24. Lawson CD, Ridley AJ. Rho GTPase signaling complexes in cell migration and invasion. J Cell Biol. 2018 Feb 5;217(2):447–57.

25. Rappel WJ, Edelstein-Keshet L. Mechanisms of Cell Polarization. Curr Opin Syst Biol. 2017 Jun;3:43–53.

26. Chiou JG, Balasubramanian MK, Lew DJ. Cell Polarity in Yeast. Annu Rev Cell Dev Biol. 2017 Oct 6;33:77–101.

27. Goryachev AB, Pokhilko AV. Dynamics of Cdc42 network embodies a Turing-type mechanism of yeast cell polarity. FEBS Lett. 2008 Apr 30;582(10):1437–43.

28. Woods B, Lew DJ. Polarity establishment by Cdc42: Key roles for positive feedback and differential mobility. Small GTPases. 2019 Mar;10(2):130.

29. Mori Y, Jilkine A, Edelstein-Keshet L. Wave-Pinning and Cell Polarity from a Bistable Reaction-Diffusion System. Biophys J. 2008 May 1;94(9):3684–97.

30. Otsuji M, Ishihara S, Co C, Kaibuchi K, Mochizuki A, Kuroda S. A Mass Conserved Reaction–Diffusion System Captures Properties of Cell Polarity. PLOS Comput Biol. 2007 Jun 8;3(6):e108.

31. Xu B, Jilkine A. Modeling the Dynamics of Cdc42 Oscillation in Fission Yeast. Biophys J. 2018 Feb 6;114(3):711–22.

32. Gerisch G, Prassler J, Butterfield N, Ecke M. Actin Waves and Dynamic Patterning of the Plasma Membrane. Yale J Biol Med. 2019 Sep 20;92(3):397–411.

33. Beta C, Edelstein-Keshet L, Gov N, Yochelis A. From actin waves to mechanism and back: How theory aids biological understanding. Michelot A, Akhmanova A, editors. eLife. 2023 Jul 10;12:e87181.

34. Negrete J, Pumir A, Westendorf C, Tarantola M, Bodenschatz E, Beta C. Receptor-induced transient responses in cells with oscillatory actin dynamics. Phys Rev Res. 2020 Mar 2;2(1):013239.

35. Marinović M, Šoštar M, Filić V, Antolović V, Weber I. Quantitative imaging of Rac1 activity in Dictyostelium cells with a fluorescently labelled GTPase-binding domain from DPAKa kinase. Histochem Cell Biol. 2016 Sep;146(3):267–79.

36. Hodge RG, Ridley AJ. Regulating Rho GTPases and their regulators. Nat Rev Mol Cell Biol. 2016 Aug;17(8):496–510.

38. Goryachev AB, Leda M. Autoactivation of small GTPases by the GEF–effector positive feedback modules. F1000Research. 2019 Sep 23;8:F1000 Faculty Rev-1676.

39. Khatibi S, Rios KI, Nguyen LK. Computational Modeling of the Dynamics of Spatiotemporal Rho GTPase Signaling: A Systematic Review. Methods Mol Biol Clifton NJ. 2018;1821:3–20.

40. Miao Y, Bhattacharya S, Banerjee T, Abubaker-Sharif B, Long Y, Inoue T, et al. Wave patterns organize cellular protrusions and control cortical dynamics. Mol Syst Biol. 2019 Mar;15(3):e8585.

41. Chang F, Lemmon C, Lietha D, Eck M, Romer L. Tyrosine Phosphorylation of Rac1: A Role in Regulation of Cell Spreading. PLOS ONE. 2011 Dec 6;6(12):e28587.

42. Bunda S, Heir P, Srikumar T, Cook JD, Burrell K, Kano Y, et al. Src promotes GTPase activity of Ras via tyrosine 32 phosphorylation. Proc Natl Acad Sci U S A. 2014 Sep 9;111(36):E3785–3794.

43. Johnson JL, Erickson JW, Cerione RA. New insights into how the Rho guanine nucleotide dissociation inhibitor regulates the interaction of Cdc42 with membranes. J Biol Chem. 2009 Aug 28;284(35):23860–71.

44. Moissoglu K, Slepchenko BM, Meller N, Horwitz AF, Schwartz MA. In Vivo Dynamics of Rac-Membrane Interactions. Mol Biol Cell. 2006 Jun;17(6):2770–9.

45. Khalili B, Lovelace HD, Rutkowski DM, Holz D, Vavylonis D. Fission Yeast Polarization: Modeling Cdc42 Oscillations, Symmetry Breaking, and Zones of Activation and Inhibition. Cells. 2020 Jul 24;9(8):1769.

46. Khalili B, Merlini L, Vincenzetti V, Martin SG, Vavylonis D. Exploration and stabilization of Ras1 mating zone: A mechanism with positive and negative feedbacks. PLOS Comput Biol. 2018 Jul 20;14(7):e1006317.

47. Jung G, Pan M, Alexander CJ, Jin T, Hammer JA. Dual regulation of the actin cytoskeleton by CARMIL-GAP. J Cell Sci. 2022 Jun 15;135(12):jcs258704.

48. Loovers HM, Kortholt A, de Groote H, Whitty L, Nussbaum RL, van Haastert PJM. Regulation of phagocytosis in Dictyostelium by the inositol 5-phosphatase OCRL homolog Dd5P4. Traffic Cph Den. 2007 May;8(5):618–28.

49. Amin E, Jaiswal M, Derewenda U, Reis K, Nouri K, Koessmeier KT, et al. Deciphering the Molecular and Functional Basis of RHOGAP Family Proteins: A SYSTEMATIC APPROACH TOWARD SELECTIVE INACTIVATION OF RHO FAMILY PROTEINS. J Biol Chem. 2016 Sep 23;291(39):20353–71.

50. Yamazaki D, Itoh T, Miki H, Takenawa T. srGAP1 regulates lamellipodial dynamics and cell migratory behavior by modulating Rac1 activity. Mol Biol Cell. 2013 Nov 1;24(21):3393–405.

51. Peng X, Wang T, Gao H, Yue X, Bian W, Mei J, et al. The interplay between IQGAP1 and small GTPases in cancer metastasis. Biomed Pharmacother Biomedecine Pharmacother. 2021 Mar;135:111243.

52. Wu CF, Lew DJ. Beyond symmetry-breaking: competition and negative feedback in GTPase regulation. Trends Cell Biol. 2013 Oct 1;23(10):476–83.

53. Howell AS, Jin M, Wu CF, Zyla TR, Elston TC, Lew DJ. Negative feedback enhances robustness in the yeast polarity establishment circuit. Cell. 2012 Apr 13;149(2):322–33.

54. Das M, Drake T, Wiley DJ, Buchwald P, Vavylonis D, Verde F. Oscillatory dynamics of Cdc42 GTPase in the control of polarized growth. Science. 2012 Jul 13;337(6091):239–43.

55. Carlin LM, Evans R, Milewicz H, Fernandes L, Matthews DR, Perani M, et al. A targeted siRNA screen identifies regulators of Cdc42 activity at the natural killer cell immunological synapse. Sci Signal. 2011 Nov 29;4(201):ra81.

56. Bement WM, Leda M, Moe AM, Kita AM, Larson ME, Golding AE, et al. Activator-inhibitor coupling between Rho signalling and actin assembly makes the cell cortex an excitable medium. Nat Cell Biol. 2015 Nov;17(11):1471–83.

57. Landino J, Leda M, Michaud A, Swider ZT, Prom M, Field CM, et al. Rho and F-actin self-organize within an artificial cell cortex. Curr Biol CB. 2021 Dec 20;31(24):5613–5621.e5.

58. Hwang JU, Gu Y, Lee YJ, Yang Z. Oscillatory ROP GTPase Activation Leads the Oscillatory Polarized Growth of Pollen Tubes. Mol Biol Cell. 2005 Nov;16(11):5385–99.

59. Killich T, Plath PJ, Wei X, Bultmann H, Rensing L, Vicker MG. The locomotion, shape and pseudopodial dynamics of unstimulated Dictyostelium cells are not random. J Cell Sci. 1993 Dec 1;106(4):1005–13.

60. Vicker MG. F-actin assembly in Dictyostelium cell locomotion and shape oscillations propagates as a self-organized reaction–diffusion wave. FEBS Lett. 2002;510(1–2):5–9.

61. Maeda YT, Inose J, Matsuo MY, Iwaya S, Sano M. Ordered Patterns of Cell Shape and Orientational Correlation during Spontaneous Cell Migration. PLOS ONE. 2008 Nov 17;3(11):e3734.

62. Yamao M, Naoki H, Kunida K, Aoki K, Matsuda M, Ishii S. Distinct predictive performance of Rac1 and Cdc42 in cell migration. Sci Rep. 2015 Dec 4;5(1):17527.

63. Arai Y, Shibata T, Matsuoka S, Sato MJ, Yanagida T, Ueda M. Self-organization of the phosphatidylinositol lipids signaling system for random cell migration. Proc Natl Acad Sci. 2010 Jul 6;107(27):12399–404.

64. Shibata T, Nishikawa M, Matsuoka S, Ueda M. Modeling the self-organized phosphatidylinositol lipid signaling system in chemotactic cells using quantitative image analysis. J Cell Sci. 2012 Nov 1;125(Pt 21):5138–50.

65. Graziano BR, Gong D, Anderson KE, Pipathsouk A, Goldberg AR, Weiner OD. A module for Rac temporal signal integration revealed with optogenetics. J Cell Biol. 2017 Aug 7;216(8):2515–31.

66. Inoue T, Meyer T. Synthetic activation of endogenous PI3K and Rac identifies an AND-gate switch for cell polarization and migration. PloS One. 2008 Aug 27;3(8):e3068.

67. Srinivasan S, Wang F, Glavas S, Ott A, Hofmann F, Aktories K, et al. Rac and Cdc42 play distinct roles in regulating PI(3,4,5)P3 and polarity during neutrophil chemotaxis. J Cell Biol. 2003 Feb 3;160(3):375–85.

68. Weiner OD, Neilsen PO, Prestwich GD, Kirschner MW, Cantley LC, Bourne HR. A PtdInsP(3)- and Rho GTPase-mediated positive feedback loop regulates neutrophil polarity. Nat Cell Biol. 2002 Jul;4(7):509–13.

69. Wigbers MC, Brauns F, Hermann T, Frey E. Pattern localization to a domain edge. Phys Rev E. 2020 Feb;101(2–1):022414.

70. Ladwein M, Rottner K. On the Rho’d: The regulation of membrane protrusions by Rho-GTPases. FEBS Lett. 2008;582(14):2066–74.

71. Pertz O. Spatio-temporal Rho GTPase signaling - where are we now? J Cell Sci. 2010 Jun 1;123(Pt 11):1841–50.

72. Marston DJ, Anderson KL, Swift MF, Rougie M, Page C, Hahn KM, et al. High Rac1 activity is functionally translated into cytosolic structures with unique nanoscale cytoskeletal architecture. Proc Natl Acad Sci U S A. 2019 Jan 22;116(4):1267–72.

73. Pollard TD. Regulation of actin filament assembly by Arp2/3 complex and formins. Annu Rev Biophys Biomol Struct. 2007;36:451–77.

74. Bolado-Carrancio A, Rukhlenko OS, Nikonova E, Tsyganov MA, Wheeler A, Garcia-Munoz A, et al. Periodic propagating waves coordinate RhoGTPase network dynamics at the leading and trailing edges during cell migration. eLife. 2020 Jul 24;9:e58165.

75. Gardiner EM, Pestonjamasp KN, Bohl BP, Chamberlain C, Hahn KM, Bokoch GM. Spatial and Temporal Analysis of Rac Activation during Live Neutrophil Chemotaxis. Curr Biol. 2002 Dec 10;12(23):2029–34.

76. Zhou S, Li P, Liu J, Liao J, Li H, Chen L, et al. Two Rac1 pools integrate the direction and coordination of collective cell migration. Nat Commun. 2022 Oct 12;13(1):6014.

77. Jilkine A, Marée AFM, Edelstein-Keshet L. Mathematical model for spatial segregation of the Rho-family GTPases based on inhibitory crosstalk. Bull Math Biol. 2007 Aug;69(6):1943–78.

78. Novák B, Tyson JJ. Design principles of biochemical oscillators. Nat Rev Mol Cell Biol. 2008 Dec;9(12):981–91.

79. Kuo CC, Savage NS, Chen H, Wu CF, Zyla TR, Lew DJ. Inhibitory GEF phosphorylation provides negative feedback in the yeast polarity circuit. Curr Biol CB. 2014 Mar 31;24(7):753–9.

80. Wu CF, Chiou JG, Minakova M, Woods B, Tsygankov D, Zyla TR, et al. eLife. eLife Sciences Publications Limited; 2015 [cited 2023 Feb 24]. Role of competition between polarity sites in establishing a unique front. Available from: https://elifesciences.org/articles/11611

81. Das M, Nuñez I, Rodriguez M, Wiley DJ, Rodriguez J, Sarkeshik A, et al. Phosphorylation-dependent inhibition of Cdc42 GEF Gef1 by 14-3-3 protein Rad24 spatially regulates Cdc42 GTPase activity and oscillatory dynamics during cell morphogenesis. Mol Biol Cell. 2015 Oct 1;26(19):3520–34.

82. Chiou JG, Ramirez SA, Elston TC, Witelski TP, Schaeffer DG, Lew DJ. Principles that govern competition or co-existence in Rho-GTPase driven polarization. PLOS Comput Biol. 2018 Apr 12;14(4):e1006095.

83. Goryachev AB, Leda M. Many roads to symmetry breaking: molecular mechanisms and theoretical models of yeast cell polarity. Mol Biol Cell. 2017 Feb 1;28(3):370–80.

84. Chiou J geng, Moran KD, Lew DJ. How cells determine the number of polarity sites. Balasubramanian MK, Barkai N, editors. eLife. 2021 Apr 26;10:e58768.

85. Jacobs B, Molenaar J, Deinum EE. Small GTPase patterning: How to stabilise cluster coexistence. PLOS ONE. 2019 Mar 7;14(3):e0213188.

86. Denk-Lobnig M, Martin AC. Modular regulation of Rho family GTPases in development. Small GTPases. 2017 Mar 17;10(2):122–9.

87. Houk AR, Jilkine A, Mejean CO, Boltyanskiy R, Dufresne ER, Angenent SB, et al. Membrane tension maintains cell polarity by confining signals to the leading edge during neutrophil migration. Cell. 2012 Jan 20;148(1–2):175–88.

88. Mierke CT, Puder S, Aermes C, Fischer T, Kunschmann T. Effect of PAK Inhibition on Cell Mechanics Depends on Rac1. Front Cell Dev Biol [Internet]. 2020 [cited 2023 Sep 7];8. Available from: https://www.frontiersin.org/articles/10.3389/fcell.2020.00013

89. Wu M, Liu J. Mechanobiology in cortical waves and oscillations. Curr Opin Cell Biol. 2021 Feb 1;68:45–54.

90. Karsenti E. Self-organization in cell biology: a brief history. Nat Rev Mol Cell Biol. 2008 Mar;9(3):255–62.

91. Kruse K, Joanny JF, Jülicher F, Prost J, Sekimoto K. Generic theory of active polar gels: a paradigm for cytoskeletal dynamics. Eur Phys J E. 2005 Jan 1;16(1):5–16.

92. Beljan S, Herak Bosnar M, Ćetković H. Rho Family of Ras-Like GTPases in Early-Branching Animals. Cells. 2020 Oct 13;9(10):2279.

93. Kimmel AR, Faix J. Generation of multiple knockout mutants using the Cre-loxP system. Methods Mol Biol Clifton NJ. 2006;346:187–99.

94. Dormann D, Libotte T, Weijer CJ, Bretschneider T. Simultaneous quantification of cell motility and protein-membrane-association using active contours. Cell Motil. 2002;52(4):221–30.

